# Data science and its future in large neuroscience collaborations

**DOI:** 10.1101/2024.03.20.585936

**Authors:** Manuel Schottdorf, Guoqiang Yu, Edgar Y. Walker

**Affiliations:** Princeton Neuroscience Institute, Princeton University, Princeton, NJ, USA; Bradley Department of Electrical and Computer Engineering, Virginia Polytechnic Institute and State University, Arlington, VA, USA; Department of Physiology and Biophysics, Computational Neuroscience Center, University of Washington, Seattle, WA, USA

## Abstract

The rise of large scientific collaborations in neuroscience requires systematic, scalable, and reliable data management. How this is best done in practice remains an open question. To address this, we conducted a data science survey among currently active U19 grants, funded through the NIH’s BRAIN Initiative. The survey was answered by both data science liaisons and Principal Investigators, speaking for ∼500 researchers across 21 nation-wide collaborations. We describe the tools, technologies, and methods currently in use, and identify several shortcomings of current data science practice. Building on this survey, we develop plans and propose policies to improve data collection, use, publication, re-use and training in the neuroscience community.

## Introduction

New kinds of data are being collected at increasing rates, challenging established methods for data processing and interpretation. In light of these changes, biology is moving towards collaborative team science, often combining experimental work with tools and instruments from statistics, physics, mathematics, engineering, and computer science^1,2^. Data science can be the key catalyst for this transition^3–6^. A recent example of this are large collaborative U19 projects, funded through the NIH’s BRAIN Initiative which seek to understand circuits of the central nervous system^4,7–10^.

From 2022 until early 2023 we conducted a survey across U19s to identify (1) tools, technologies, and methods currently in use across these collaborations, and (2) shared difficulties and challenges to data science that could best be addressed by a community-wide effort. In the first part of this article, we will summarize key insights from the survey, and in the second part present proposals to accelerate scientific progress by improving (1) data integration, (2) data and code sharing, and (3) training and best practices for workforce development. We hope that this work can improve scientific standards in our community, and the life sciences more broadly.

## Results

An overview of active U19s is shown in **Fig. 1A**. Our survey was distributed among these collaborations, every one of which features a data science core to manage analysis, archiving and data sharing. The survey was answered by both data science liaisons managing these cores, and Principal Investigators (PIs). The survey contained six sections focused on (1) team composition, (2) analytical and software tools currently in use, (3) software engineering, standards, and data/code sharing, (4) Personnel and budget, (5) Challenges, and (6) Future Steps totaling 82 questions. From N=21 currently active U19s, n=15 U19s provided us with the data presented here, corresponding to an answer rate of 71%. The missing U19s were ignored in our analysis. We speculate, in the discussion section, on why no replies could be obtained. In the following paragraphs we present an overview and key insights from our survey, and actionable items to improve data science. The complete survey and analysis are provided in the supplemental material.

**Fig. 1:**
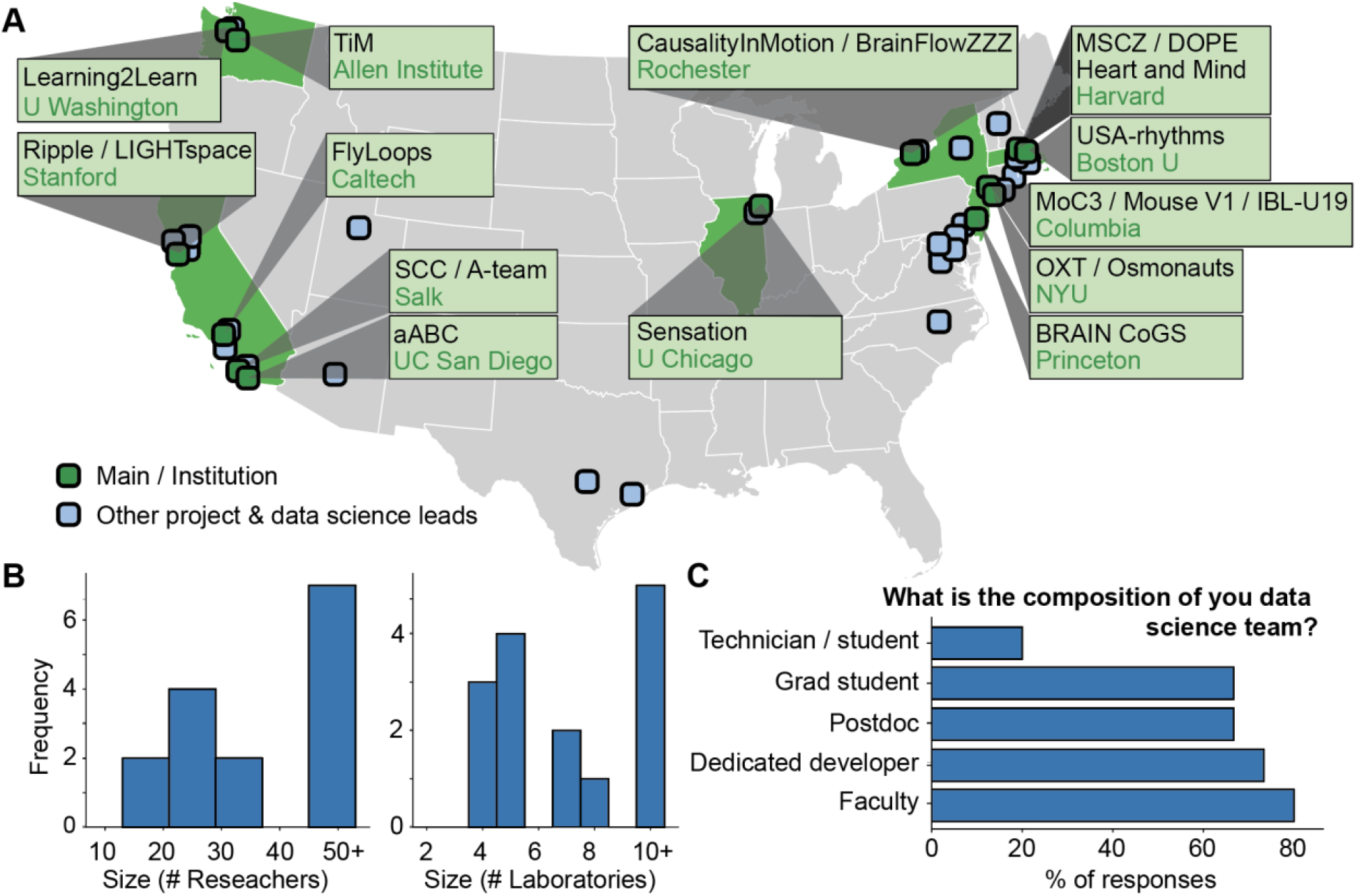
Overview of the Brain Initiative’s U19 program. A) Title and locations of N=21 currently active U19s across the nation (green boxes) and affiliation of individual projects as well as datacore investigators (blue boxes). B) Team size of the U19s and number of participating laboratories. A typical U19 team has ∼40 scientists across ∼7 labs. This survey covers ∼500 researchers. C) Composition and role of data science core personnel. Notice that most data science is done by postdocs, developers, or faculty suggesting advanced training requirements.

While the U19 program funds nationwide research, it is highly concentrated on a small subset of universities. Using biographical information of n=176 PIs and data science leads across the N=21 grants, we found that 26.2% were affiliated with either Harvard or Columbia University. Summing over the top six institutions (Stanford, Princeton, NYU and the University of Washington) accounted for more than half of all senior personnel (50.6%) (**Fig. 1A**). The largest contributing public university was the University of Washington at 5.1%. Centered around these locations, a typical U19 team has ∼40 scientists across ∼7 labs (**Fig. 1B)** and is supported by 1–2 full time data scientists and many other members with diverse backgrounds / roles. In practice, we found that data science is often done by more senior researcher positions such as postdocs, faculty, or specific developers hired for the job (**Fig. 1C**). This suggests relatively advanced training requirements when compared to a typical research position. To test this hypothesis, we also surveyed the role of data science and job profiles.

This produced a formidable list of tasks: the development of (semi-) automatic data processing pipelines, maintenance of data infrastructure, general tech support and education for researchers, help for researchers when writing analyses Code (e.g. replication of published algorithms), support in scientific data analyses and IT resource usages, custom web GUI/frontend development, support in experimental setup and hardware interfacing, and support in integrating theory and experiment (see supplemental material for the full questionnaire). We discuss the implications of this diverse job profile further below, but it should be noted that in the private sector, most of these roles would be filled by specifically trained specialists. In academia, recruitment seems to occur on higher levels to find sufficiently well-trained individuals who can fill these very diverse roles (cf. Fig. 1C). While these roles place data science into an advantageous position to serve as a key catalyst to accelerate science, this diversity also suggests that management of such diverse roles is challenging, and data science can become prohibitively expensive. Funding is an important point that we will discuss below.

One of the specific contributions of data science is to prepare and facilitate data sharing both within collaborations and across the community. How is data shared? In practice, most data is copied through local networks or exchanged via eMail or Slack. Little data is shared with data management tools (**Fig. 2A**). When asked about such specific tools, like Neurodata without Borders (NWB), a data format based on hdf5 that is becoming a standard for neurophysiological data^11^, we discovered that these are well known and used in a majority of collaborations. However, only a small fraction of data is maintained in these formats (**Fig. 2B**). Insofar as data pipelines exist, their underlying code is generally shared. This is different for data. Even after publication, current U19s share only a small fraction of data underlying their publications. It should be noted that the distribution is bimodal, with few collaborations sharing all their data (**Fig. 2C**), while the majority of U19s only share a small fraction, if any. Even though U19 collaborations develop and publish pipelines for electrophysiology^12^ or Ca^2+^ imaging^13^ no visible standards have emerged as to what data to share. The split is almost equal between publishing raw data itself (such as tiff-stacks, or e-Phys voltage traces), Intermediate data such as Ca^2+^ transients or spike times and analyzed data such as peri-stimulus time histograms, or dimension reduced data (**Fig. 2D**). Among the individually developed software, at most ∼50% of critical infrastructure code has gone through review or pull requests (**Fig. 2E**). We will discuss the implications of these findings below.

**Fig. 2:**
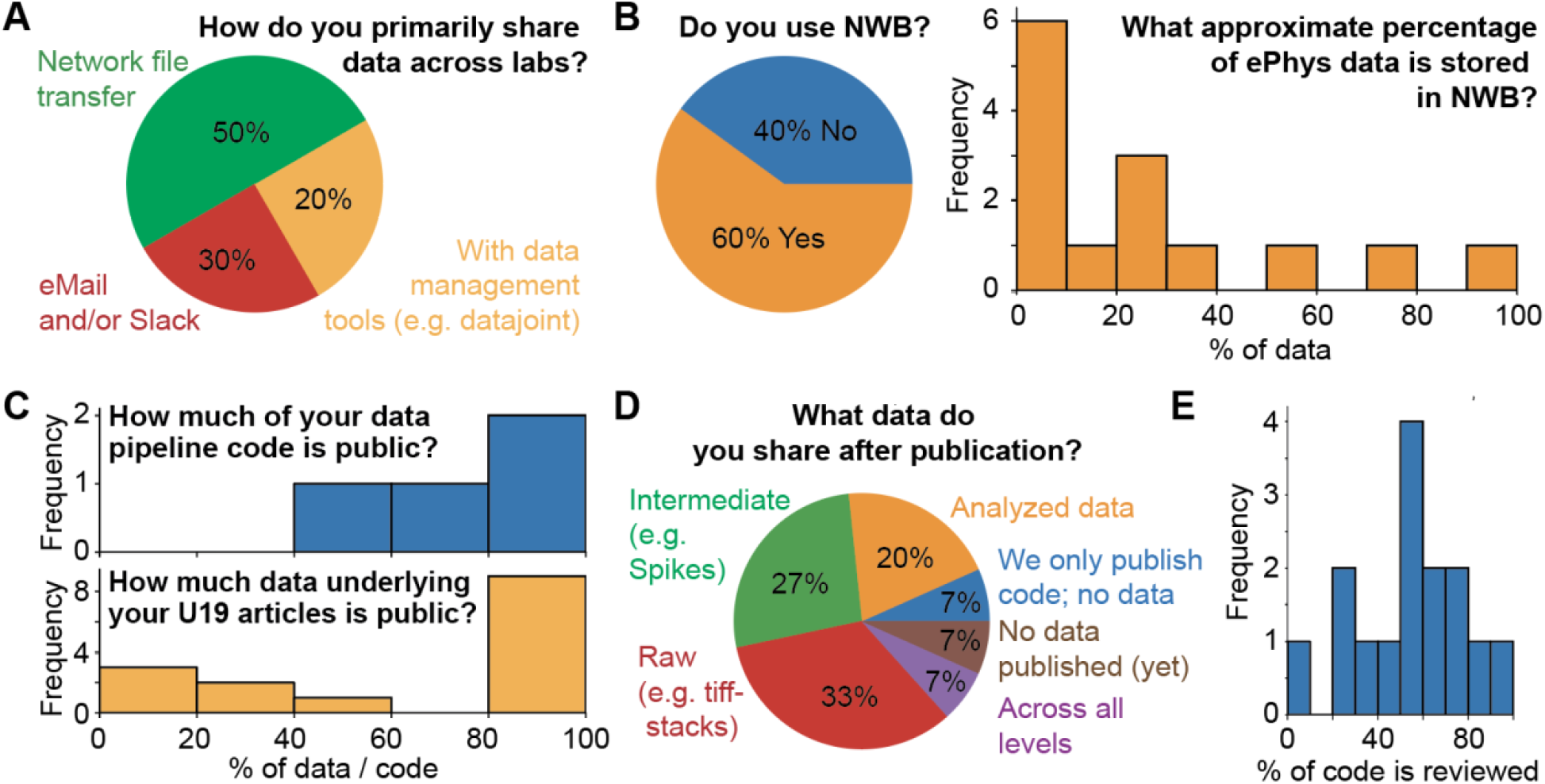
Key results of current data sharing practices. A) The main mechanisms by which data are exchanged are Slack/eMails and Network transfer. Not much is standardized. B) Most U19s are aware of, and use tools like Neurodata without Borders (NWB). But only in a limited capacity. C) Much code is shared, but data sharing is bimodal. D) Data is being published at all stages of processing without clear structure. E) A significant fraction of code has never been reviewed.

Is limited training a key factor for the limited adoption of better data science practices? We found that theory and experimental researchers are similarly interested and engaged in data science training. However, only around 50% of experimental researchers have the necessary coding skills to implement a simple algorithm (**Fig. 3A**). Theory researchers tend to be more experienced regarding coding and data science but are further away from experimental data collection and processing pipelines. While many U19s provide training for their researchers, it is limited in scope and often restricted to onboarding documents (**Fig. 3B**). Only ∼50% of collaborations enforce standards for data science practices. The rest leaves data science practices up to the best (but limited) abilities of the respective researcher (**Fig. 3C**). The limited training and lack of standards in most U19 collaborations likely contributes to the limited adoption of better data science practices.

**Fig. 3:**
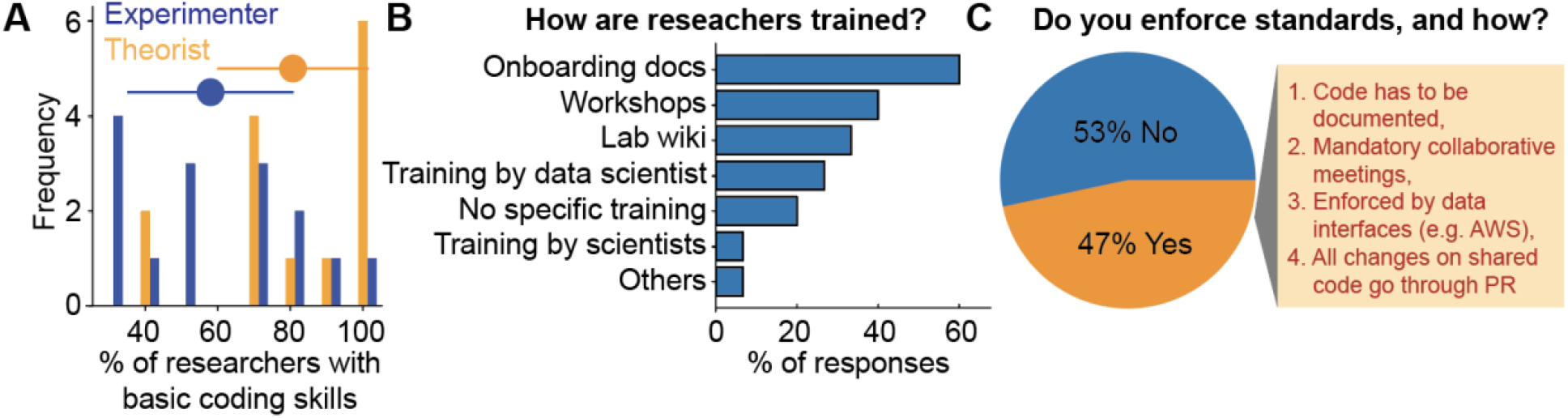
Key results of researcher qualifications, training, and management. A) Fraction of theory and experimental researchers that can implement a simple algorithm. Theory researchers are typically more competent, but only around ∼50% of experimental researchers have basic coding experience. The precise question was “*What fraction of your U19’s experimental researchers has reasonable coding experience? (e*.*g. can implement a simple algorithm like bubble sort in python, or use the slurm engine on a cluster*.)“ Dots indicate means and C.I. B) Current training is done, but limited in scope. C) Most U19s do not enforce standards. The ones who do enforce standards do so on a superficial level.

## Actions and policies to improve data science

We believe the current state of the field, documented above, necessitates improvements. To aid this process we presented our results to the U19 data science consortium in January 2023, conducted a workshop at the 2023 Brain initiative meeting in Washington D.C. in June, and spoke at a SfN Symposium in November 2023. In the following paragraphs, we summarize strategies and proposals discussed at these events.

### Integration

Neuroscientists are seeking increasingly close collaborations with experts in computation, statistics, and theory to mine and understand their data. These interactions and associated analysis methods, many of which did not exist 10 years ago, force us to reconceive what it means to be an experimental neuroscientist today. We believe that this scientist of the future has deep knowledge in one discipline and requires basic fluency in several others, including data science^14^. We consider improvements in workforce training below. Here, we list policies that can aid this integration directly through three complementary strategies: *<u>First</u>*, new funding schemes could incentivize better coding and data sharing practices. For example, expertise could be recruited from other scientific fields into Neuroscience via career transition grants, or grants to maintain software beyond the lifetime of an individual U19. While the former exists in some private foundations, the NIH has supported the latter through a NOSI in 2022^15^. These efforts should be scaled up. <u>Second</u>, data scientists should be included early during data collection to foster a spirit of collaboration rather than perceived competition between data collection and processing^2^. They should also be present in lab meetings to become a normal part of any laboratory involved with collecting large neurophysiological data. Without this collaborative spirit, and clear roles, experimentalists might feel devalued to a technical role limited to data collection, and do not see the advantages of outsourcing coding to an expert. *<u>Third</u>*, a specific idea to better integrate U19s is the generation of *virtual* or *online* data core facilities that are open to researchers across the nation. This distributed core facility would be particularly suited to define and enforce standards in the field.

### Sharing

We found that across U19s, there is a shared interest to improve sharing and standardization of data formats, analysis, processing code, data pipelines and data infrastructures, but with limited success. Here we present four suggestions to ameliorate our standards: *<u>First</u>*, laboratories should use well-established code whenever possible. We are hopeful that with better training on new methods, and improved communication (see below) much of this can be addressed easily, precisely because there is a shared sense of urgency and usefulness. Making the reuse of code easier is the best motivation for researchers to not develop from scratch and to not reinvent what already exists. *<u>Second</u>*, we have to establish common guidelines for code and data sharing combined with documentation. This requires that we agree on how data should be shared, and much work has already been accomplished in solving this issue^16–18^. It also requires that we agree on what data should be shared. We advocate for sharing of both: (1) raw data to aid future discoveries, and (2) pre-processed data and associated code to aid reproducibility efforts and quality of science. This is particularly important if custom pre-processing pipelines are in use. We should hold people accountable if this data is not provided. This could be enforced by the NIH itself, for example by providing *Institutional Certifications* for minimum institutional standards or by making the most recent NOT-OD-21-013 even stronger. It can also be enforced by us as a community. *<u>Third</u>*, as a community, also on an international scale^5^, we can decide that all papers reviewed *by us* need to publish code and data. The authors of this article have adopted this stance as reviewers. Along these lines, and to advertise for a growing body of published datasets, we wish to highlight datasets across scales that have already been published to the highest standards. Key examples are work published by the Allen Institute^18,19^ and IBL^20,21^. *<u>Fourth</u>*, we propose to generate public infrastructure outside of the Amazon/Microsoft/Google ecosystem. In other fields of science, this has been an enormous success, considering databases like GenBank, or pubmed itself. A public repository will have the benefits of (1) being cheaper on the taxpayer as not ultimately operating for profit and (2) access as a .gov domain will make data-publications citable in NIH grants.

### Training

We need to better train researchers. This point came up several times in the paragraphs above. *<u>First</u>*, we propose that training grants, like the NIH’s T32 mechanism, have to contain at least one introductory coding class, and one course on computational methods. We wish to stress that any study program that allows students to graduate without mandatory and formal training to code is doing these students a grave disservice for their individual careers and the progress of science. It should also be noted that these are among the skills that are directly transferable to an industry setting. Acknowledging that not all students and postdocs will obtain faculty positions, we should do our students this service. There was also a remarkable consensus in our workshop that it is easier to train a computer scientist to perform rodent survival surgeries than to train a biologist to set up a relational database with a simple interface. Classwork can address this. *<u>Second</u>*, we should provide training opportunities for using tools and adopting best practices for existing staff, possibly combined with grants that allow us to recruit such expertise, which is readily available in math, physics, computer science, and engineering. This could happen across levels, using a F99/K00 mechanism to recruit postdocs, or K99/R00 to recruit junior faculty specifically with degrees in these underrepresented scientific fields. If a lab is already bad at data science, training within this lab is pointless. Only recruiting external expertise ameliorates this. Once recruited, we propose to pair up postdocs and students into mentor/mentee teams in which knowledge transfer can happen effectively and informally. Researchers experienced in data science can be enormously beneficial for guiding students or beginning postdocs. Researchers with expertise in mouse behavior and neurophysiology could ideally be teamed up with someone experienced in modeling or population level analysis so that the overall quality of the project improves, as well as an improved training experience can be provided for the individual researchers. *<u>Third</u>*, while it would be more effective to train students in earlier career stages, we are faced with the reality of limited computational abilities, in particular by postdoctoral researchers. To address this practically, and on short time scales, we should establish shared/common training material across our U19s and set up regular seminars or webinars covering common interests across U19. This could complement modern large scale online resources like the Neuromatch Academy^22^, or stand on its own.

## Discussion

With increasingly powerful techniques come new data sets of massive size and complexity. These big data sets can radically accelerate the BRAIN Initiative^4^ and are accumulating at an unprecedented rate. This will accelerate as the BRAIN Initiative gathers further momentum. The primary tools to address these data are robust data infrastructure and good data science. To survey the current state of data science, we conducted a comprehensive survey across BRAIN initiative funded U19 projects. We then identified ways how data science can be improved.

We ran our study over several months via google surveys. This allowed us to continuously track submissions, and easily send reminders. Despite our best efforts and multiple reminders, ∼25% of U19s did not fill in our survey. While we have no specific insight as to why these respective data science liaisons and PIs chose to ignore multiple requests to contribute to this work, personal communication in two instances revealed that one researcher was a new postdoc who felt not familiar enough with the scope and program of their U19s to answer the survey. A second researcher was on parental leave, and their position was vacant in the meantime. While both examples were somewhat concerning for their respective U19s, this suggests that leaving out these U19 has not produced systematic distortions of what was represented here, as reasons were individual and not related to data science itself.

In the vast majority of U19s, coding practices and data management were limited and significantly lagged behind scientific standards in other fields (e.g.^23^), and industry and also open-source project (e.g.^24^). Even highly popular tools across U19s, like kilosort and spikeGLX, were written by single individuals without code review. This is surprising as many U19 researchers are in fact involved in other open-source projects and are aware of best practices. For example, we ourselves have been involved in^21,25,26^. Bringing this knowledge into our respective U19s is often challenging because of the complexity and reality of data science in collaborations where most workers have limited computational abilities. Consistent with this, we found that at present google documents and Slack play a much larger role to distribute and organize data than *any* established tool in either the scientific or private sector. We hope that this article can raise awareness that this is concerning and will need to change in the future.

Data science funding might be a critical bottleneck. We discovered that the required job profile of a typical U19 data scientist is highly diverse. This diversity will require experienced workers, necessarily at a higher pay scale than traditional postdocs. While we can possibly retain workers with a pay-cut of ∼50% relative to an industry position, the current split exceeds 300%. We believe this is not sustainable. To this end we have proposed alternative means of addressing this in the *Actions* section, for example by recruiting young early-career researchers with data science experience into the life sciences with specific funding programs.

We are not the first community to walk this path. Data repositories for curating and providing access to scientific data, combined with preserving analysis workflows in many domains became a driving force for progress in both industry and academia. <u>First</u>, considerable work was done on standardization in the industry, for example through norms DIN31644^27^ and ISO16363^28^. Our colleagues from economics have also estimated the cost and benefits of digitization with the advent of digital computers in the 1970s and 1980s. Empirically, for the first ∼15 years, the use of digital computers *slowed* growth in productivity, a phenomenon known as the *productivity paradox*, before harnessing the enormous long-term benefits of digitization. This phenomenon arose because businesses had to invest in technology and learn how to use this technology before integrating them into core processes and business models^29,30^. It is conceivable that similar effects will play out in our communities, even though our hope is to make this transition significantly faster. <u>Second</u>, in science, desiderata for data repositories were already articulated via FAIR (Findable, Accessible, Interoperable and Reusable)^31^ and more recently the TRUST principles (Transparency, Responsibility, User focus, Sustainability, Technology) for digital repositories^32^. Data repositories will also allow to measure data impact^33,34^, for example by tracking download and citation counts^35^ and further the use and utility data^36,37^. In specific scientific fields this progress is happening rapidly. This is exemplified in cognitive science^38,39^ which was driven by both a stick (the replication crisis), and the carrot. For example, new fMRI processing pipelines have only few exposed parameters, and are sufficiently well documented for setup by University IT, and not an individual researcher.^40^ Raw data is easy to store on OpenNeuro^41^, and these pipelines make use of (1) established deployment instruments like docker and singularity, (2) established unit-tests to maintain functionality, and (3) code reviews to catch bugs. This ease of adoption significantly lowers initial costs of adoption.

Across life sciences, an ongoing effort over the last 50 years made data and methods more reproducible and easier to access. This reflects both enormous successes as well as failures. While a failure was mentioned above, key fields that have benefitted from good data science are structural biology^42^ and genomics^43^. GenBank, for example, contains nucleotide sequences and protein translations publicly maintained by the National Center for Biotechnology Information (NCBI; part of NIH)^44^. The systematic collection and publication of protein sequences and structures over half a century lead to scientific breakthroughs with methods that were unthinkable when the first data began to be collected, such as AlphaFold^45^. We are optimistic that similar work in neuroscience will transform the field in ways that are impossible to predict now. To this end, the establishment of public, integrated repositories for datasets and data analysis tools, with an emphasis on ready accessibility and effective central maintenance, will have immense value for the science of the future^33,46,47^.

## Conclusions

Good data science can accelerate the progress of science in predictable, and sometimes unpredictable ways. Our survey of the current state across large U19-funded neuroscience collaborations revealed substantial challenges to data science that concern us all. Only as a community can we address these challenges. Let’s get to it.

## Acknowledgements

We would like to thank the many community members who contributed to this work, including Daniela Witten and Kanaka Rajan from the BRAIN Initiative’s Theories, Models and Methods Program. And Grace C.Y. Peng, Hermon R. Gebrehiwet, Susan N. Wright, James W. Gnadt, Joseph D. Monaco, and the NIH BRAIN Initiative U19 Brain Circuits Program. MS is supported by NIH grant U19NS132720, a C.V. Starr fellowship, and a Burroughs Wellcome Fund’s Career Award at the Scientific Interface. GY is supported by NIH grant U19NS123719. EYW is supported by NIH grant U19NS107609.

